# Deep Ensemble Model for Quantitative Optical Property and Chromophore Concentration Images of Biological Tissues

**DOI:** 10.1101/2024.12.04.626749

**Authors:** Bingbao Yan, Bowen Song, Chang Ge, Xinman Yin, Wenchao Jia, Gen Mu, Yanyu Zhao

## Abstract

The ability to quantify widefield tissue optical properties (OPs, i.e., absorption and scattering) has major implications on the characterization of various physiological and disease processes. However, conventional image processing methods for tissue optical properties are either limited to qualitative analysis, or have tradeoffs in speed and accuracy. The key to quantification of optical properties is the extraction of amplitude maps from reflectance images under sinusoidal illumination of different spatial frequencies. Conventional three-phase demodulation (TPD) method has been demonstrated for the mapping of OPs, but it requires as many as 14 measurement images for accurate OP extraction, which leads to limited throughput and hinders practical translation. Although single-phase demodulation (SPD) method has been proposed to map OPs with a single measurement image, it is typically subject to image artifacts and decreased measurement accuracy. To tackle those challenges, here we develop a deep ensemble model (DEM) that can map tissue optical properties with high accuracy in a single snapshot, increasing the measurement speed by 14× compared to conventional TPD method. The proposed method was validated with measurements on an array of optical phantoms, *ex vivo* tissues, and *in vivo* tissues. The errors for OP extraction were 0.83 ± 5.0% for absorption and 0.40 ± 1.9% for reduced scattering, dramatically lower than that of the state-of-the-art SPD method (2.5 ± 15% for absorption and -1.2 ± 11% for reduced scattering). It was further demonstrated that while trained with data from a single wavelength, the DEM can be directly applied to other wavelengths and effectively obtain optical property and chromophore concentration images of biological tissues. Together, these results highlight the potential of DEM to enable new capabilities for quantitative monitoring of tissue physiological and disease processes.

## I. Introduction

The ability to quantify widefield optical properties (i.e., absorption and scattering) of strongly scattering media such as biological tissues, has major implications on the characterization of various physiological and disease processes [1], [2], [3], [4]. With quantified tissue absorption at different wavelengths, concentrations of functional tissue components, such as oxyhemoglobin and deoxyhemoglobin, can be extracted. Alterations in the concentrations and spatial distribution of these components are hallmarks of many conditions including cardiovascular disease [5], [6], tissue metabolism [7], oxygenation [8], [9], skin burns [10], [11], [12], [13], diabetes [14], [15], and several cancers [2], [16], [17]. [18], [19].

The key to quantification of tissue optical properties is the extraction of amplitude maps from reflectance images under sinusoidal illumination of different spatial frequencies (*f*_*x*_). With extracted amplitude maps, optical absorption and scattering can be quantified using a standard lookup table (LUT) generated by Monte Carlo simulation [20]. In order to get highly accurate optical property maps at a single wavelength, conventional methods utilizing three-phase demodulation (TPD) require 14 measurement images, which could lead to motion artifacts for *in vivo* measurements [21], [22]. Single-shot methods such as single-phase demodulation (SPD) have also been developed to map tissue optical properties, but these methods are typically subject to image artifacts and decreased measurement accuracy [23], [24].

To tackle those challenges, we develop a deep ensemble model (DEM) for accurate extraction of optical properties in a single snapshot. Instead of sequential image collection of different phases and spatial frequencies, our method utilizes frequency multiplexing for illumination and captures a single reflectance image that contains information of multiple spatial frequencies. The deep ensemble model is able to effectively recover demodulated (amplitude) images for each spatial frequency, from which the optical properties can be extracted.

In the remaining of this work, we first demonstrate the frequency multiplexing for the illumination pattern, which can probe tissue information of multiple spatial frequencies in a single snapshot. We then develop the deep ensemble model to effectively extract demodulated images of those spatial frequencies from the single reflectance image. We also validate the proposed DEM against conventional TPD and SPD methods through experiments on an array of optical phantoms with a wide range of optical properties as well as *ex vivo* tissues and *in vivo* tissues. While the DEM is trained with data from a single wavelength, we demonstrate that it can be directly applied to other wavelengths and effectively extract optical properties as well as concentration of important biological chromophores including oxyhemoglobin and deoxyhemoglobin. We also demonstrate the proposed method with a cuff occlusion study. To the best of our knowledge, this is the first demonstration of highly accurate mapping of tissue optical properties in a single snapshot. We conclude with a discussion of application areas in which the proposed method may have a substantial impact.

## II. Related Work

### A. Three-Phase Demodulation Method

The details of TPD method have been described elsewhere [25], [26]. Briefly, the tissue reflectance images of different phases and spatial frequencies are demodulated using Eq. (1), where *I*_1_, *I*_2_, and *I*_3_ denote the reflectance images of three different phases (0°, 120°, and 240°) with the same spatial frequency, and *M*_*ac*_ represents the demodulated image. For calibration, the diffuse reflectance image is calculated using Eq. (2), where *M*_*ac*_*tissue*_, *R*_*d*_*tissue*_, *M*_*ac*_*phantom*_ and *R*_*d*_*phantom*_ represent the demodulated image of the tissue, the diffuse reflectance image of the tissue, the demodulated image of the calibration phantom, and the diffuse reflectance image of the calibration phantom, respectively. After demodulation and calibration, the tissue diffuse reflectance images are fed into an inverse model which maps diffuse reflectance to optical properties [20], [22], [27], [28]. With optical absorption at multiple wavelengths, concentrations of endogenous chromophores, such as oxyhemoglobin and deoxyhemoglobin, can be extracted using Beer-Lambert law [4], [29].

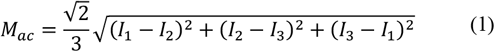

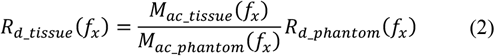

Typically, TPD method requires 5 spatial frequencies (such as [0, 0.05, 0.1, 0.2, 0.4] mm^-1^) to quantify optical properties with high accuracy [21], [22]. Since there are 3 images for each spatial frequency, it requires 14 measurement images for OP quantification (i.e., 2 images for DC and 3 images for each AC), which severely limits imaging throughput and hinders practical translation.

### B. Single-Phase Demodulation Method

To reduce the required number of images in TPD measurements, SPD methods utilize a single-phase illumination pattern and can extract both DC and AC images for OP estimation from a single measurement image. Specifically, for the demodulation, an FFT is performed on the collected image, and DC and AC components are separated by filtering in the Fourier domain [30]. The DC image is extracted from inverse FFT of the DC component, while the AC image is obtained through Hilbert transform of the AC component. Recently, Aguénounon et al. developed a convolutional neural network for the extraction of DC and AC images, resulting in reduced artifacts in demodulation compared to those obtained by filtering in the Fourier domain [23]. However, SPD methods are limited to 2 spatial frequencies for OP estimation, which has larger measurement uncertainties compared to TPD method with 5 spatial frequencies.

## III. Methods

### A. Deep Ensemble Model

To tackle the abovementioned issues, we develop a new image processing method, Deep Ensemble Model, or DEM, for the extraction of optical properties with multiple spatial frequencies (i.e., [0, 0.05, 0.1, 0.2, 0.4] mm^-1^) in a single snapshot. The 5 spatial frequencies are multiplexed and simultaneously integrated into a single illumination pattern. The demodulated images of different spatial frequencies are effectively extracted using an ensemble model based on deep learning. With DEM, the number of measurement spatial frequencies is increased from 2 to 5 compared to SPD methods (leading to significantly improved measurement accuracy), and the acquisition speed is improved by 14× compared to the TPD.

Fig. 1(a) shows the synthesis of multi-frequency pattern by frequency multiplexing, corresponding measurement images of optical phantom and hand, and their 2D Fourier spectra. For the synthesis of multi-frequency pattern, single-frequency patterns of 0.05, 0.1, 0.2, and 0.4 mm^-1^ were first rotated counter-clockwise by 0°, 25°, 50°, and 75°, respectively. After normalization, the four rotated patterns were combined to produce the multi-frequency pattern. To enhance signal-to-noise ratio of higher spatial frequency components, combination weights of 1, 1, 2, and 2 were assigned to the 0.05, 0.1, 0.2, and 0.4 mm^-1^ patterns, respectively. As shown in Fig. 1(a), the frequency components of different spatial frequencies are shifted to different locations in the Fourier domain due to different rotation angles in the multiplexing. In addition, with phantom measurement, the frequency components are visually apparent in the Fourier domain. In contrast, with hand measurement, the frequency components corresponding to the spatial frequencies are less discernible, potentially due to the existence of higher frequency contents of the object.

**Fig. 1.**
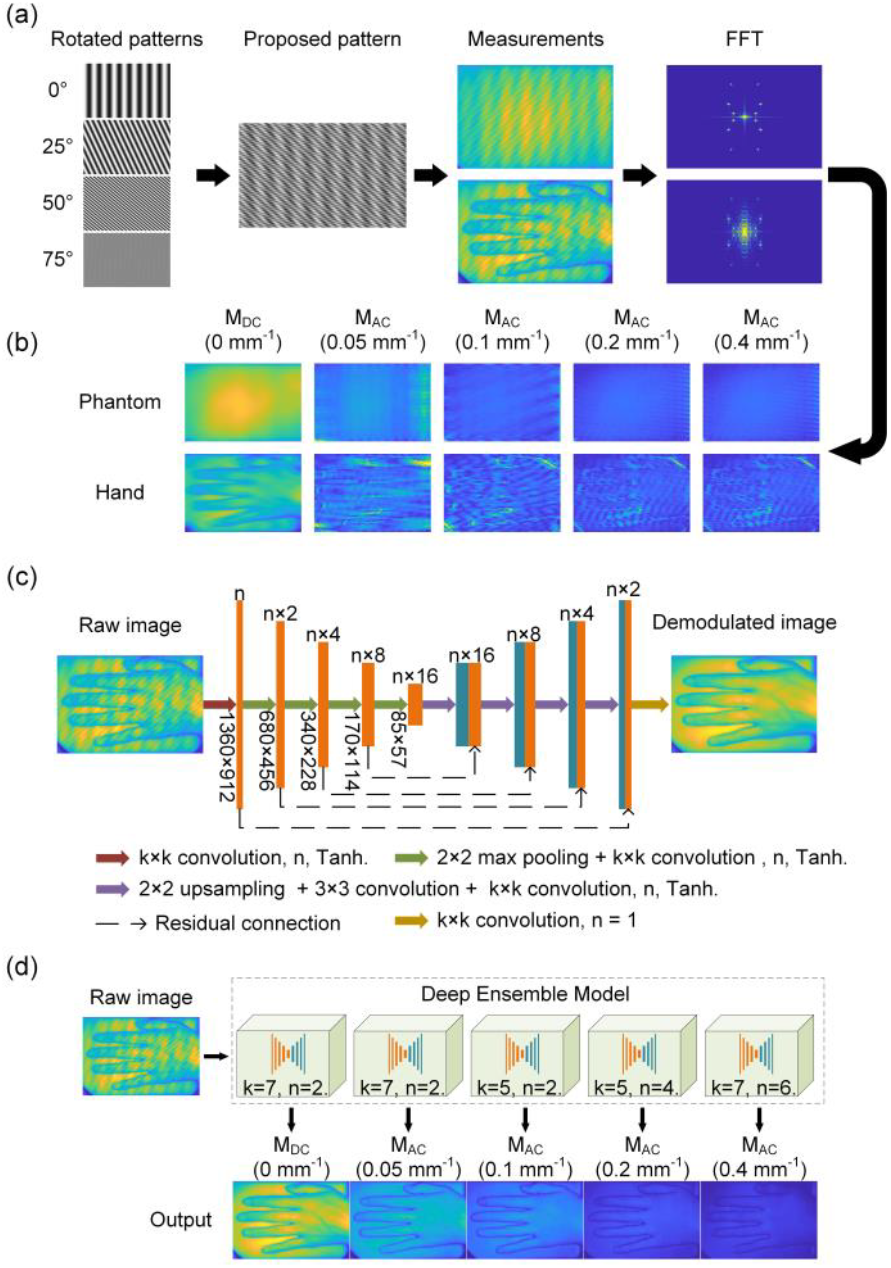
Schematics of the proposed method. (a) Synthesis of the multi-*f*_*x*_ illumination pattern, corresponding measurement images and their 2D Fourier spectra. (b) Demodulation results based on filtering in the Fourier domain with reference to SPD method. (c) CNN for the demodulation of each spatial frequency. (d) Proposed deep ensemble model and the corresponding demodulation results for 5 spatial frequencies.

For the demodulation of the 5 spatial frequencies, one may first attempt applying filters in the Fourier domain and then conducting inverse Fourier transform with reference to previous SPD methods [30]. Correspondingly, Fig. 1(b) shows demodulated images of the 5 spatial frequencies obtain by filtering in the Fourier domain, which unfortunately have artifacts for both measurements. The artifacts in the hand images appear to be more severe, potentially due to less discernible frequency components in the Fourier domain.

To reduce those artifacts, a deep ensemble model was proposed in this work. Specifically, a convolutional neural network (CNN) was first developed for the demodulation of each spatial frequency. Fig. 1(c) shows the structure of the CNN for each spatial frequency, where k and n parameters are varied for different CNNs, and represent the size and number of convolutional kernels, respectively. The CNN structure was designed with reference to U-Net, including both downsampling and upsampling operations, as well as residual connections [31]. For each CNN, the input is the measurement image encompassing 5 spatial frequencies, and the output is the demodulated image of the corresponding spatial frequency. Furthermore, an ensemble model was assembled using the 5 CNNs (each separately trained) to obtain demodulated images of the 5 spatial frequencies. The model training was performed using Python and TensorFlow framework. Mean Squared Error (MSE) was used as the loss function. The learning rate was initialized at 0.001. The Adam optimizer was chosen for optimization [32]. The training was conducted on a desktop computer with NVIDIA RTX 3090 GPU. Each CNN went through 1500 training epochs, taking approximately 5 hours. The training set consisted of 449 image sets collected at 745 nm wavelength, with image size of 1360 × 912 pixels. Among these, 399 image sets were used for training, while 50 image sets were used for validation during the training process. The loss curves for both training and validation sets had similar trends and values, suggesting no signs of overfitting in training.

As shown in Fig. 1(d), the input of the deep ensemble model is the single measurement image containing different spatial frequencies, and the output are the demodulated images of the 5 spatial frequencies, where little artifacts can be observed compared to those in Fig. 1(b). With demodulated images for the 5 spatial frequencies, optical properties can be extracted using either nearest search or a fully-connected neural network as inverse model. In this work, since the numerical range of OPs is outside that of the published fully-connected neural network, we utilized the nearest search method for OP extraction [22].

### B. Optical Instrumentation

In this study, we built a hardware system for experimental validation of the proposed method, as shown in Fig. 2. The system light source consists of three Light Emitting Diodes (LEDs) with wavelengths of 655 nm, 745 nm, and 850 nm, respectively. The output light is condensed through an aspheric lens (ACL2520U, Thorlabs, New Jersey, USA) and then directed onto a digital micromirror device (DMD, V-650L, ViALUX, Saxony, Germany) for spatial modulation. The projection lens (#33-923, Edmund Optics, New Jersey, USA) has a diameter of 30 mm and a focal length of 40 mm, and projects the modulated pattern from the DMD onto the sample. Subsequently, the reflectance image from the sample is collected using a camera (FL-20BW, Tucsen, Fuzhou, China). At a working distance of 40 cm, the field-of-view (FOV) is approximately 18 × 11 cm. To eliminate specular reflection from the sample surface, two linear polarizers (#33-083, Edmund Optics, New Jersey) are placed orthogonally in the light path. The camera captures image data with 16-bit depth and size of 1368 × 912 pixels (after 4 × 4 binning). During the acquisition, the light source, DMD, and camera are synchronized using transistor-transistor logic (TTL) signals generated by an Arduino board. Data processing is conducted on the desktop workstation equipped with an NVIDIA RTX 3090 GPU, 64 GB RAM, and an Intel Xeon® Silver 4210 R CPU.

**Fig. 2.**
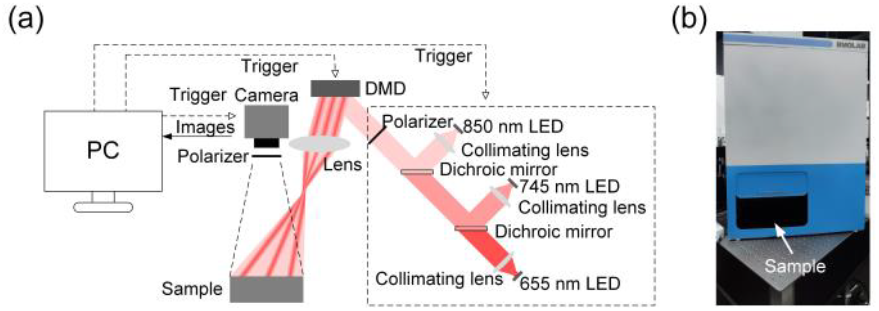
(a) System diagram. (b) Photo of the system.

### C. Data Acquisition

Data collection was conducted on optical phantoms, *ex vivo* pork tissues (including liver, muscle, stomach, intestine, and esophagus), and *in vivo* tissue (human hands), using [655, 745, 850] nm wavelengths and [0, 0.05, 0.1, 0.2, 0.3, 0.4] mm^-1^ spatial frequencies. The choice of spatial frequencies for data collection was with reference to literature [21], [22]. Specifically, the [0, 0.05, 0.1, 0.2, 0.4] mm^-1^ spatial frequencies were used for multi-*f*_*x*_ (i.e., 5-*f*_*x*_) TPD method, and the [0, 0.1, 0.2, 0.3, 0.4] mm^-1^ spatial frequencies were used for SPD method with different *f*_*x*_ combinations. For each sample, one set of measurement images was collected for six spatial frequencies (i.e., [0, 0.05, 0.1, 0.2, 0.3, 0.4] mm^-1^) with three phases, and for the proposed multi-frequency pattern. Consequently, each measurement set had 18 images (2 images for DC, 3 images for each AC, and 1 image for the multi-*f*_*x*_ pattern). The collected images, with size of 1368 × 912 pixels, were cropped to 1360 × 912 pixels (removing edge columns) to match the required data dimension and feed into the deep learning models.

The training dataset had 449 measurement sets, consisting of 204 image sets from 70 liquid phantoms measured with different phantom locations, 26 image sets from 13 solid phantoms measured with different orientations and locations, 100 image sets from 34 *ex vivo* pork tissues (including liver, muscle, stomach, intestine, and esophagus) measured with different combinations and orientations, and 119 image sets from 12 different human hands (two female subjects and four male subjects) measured with various orientations. In addition, the experimental procedures were reviewed and approved by the Beihang University Biological and Medical Ethics Committee. Moreover, the liquid phantoms were made of water, nigrosin, and intralipid. The amount of nigrosin and intralipid were varied to change the optical properties. The solid phantoms were fabricated with silicone base, and varied with different amounts of titanium dioxide and nigrosin for different optical properties.

The measurement data of 745 nm wavelength was used for training the deep learning models. With the training dataset, 399 image sets were designated for training, while the remaining 50 image sets were used for validation. The OP range of training dataset were [0.0040, 0.22] mm^-1^ for absorption and [0.30, 3.9] mm^-1^ for reduced scattering, respectively.

Moreover, test dataset was collected with the three wavelengths from additional samples different from those in the training dataset. The test dataset consists of 75 measurement sets, including 49 image sets of liquid phantoms, 18 image sets of *ex vivo* pork tissues, and 8 image sets of human hands (two female subjects and two male subjects). Additionally, the OP range of test set with three wavelengths were [0.0020, 0.33] mm^-1^ for absorption and [0.28, 4.4] mm^-1^ for reduced scattering, respectively, covering a larger range of OPs compared to the training set.

### D. Experimental Validation

The proposed DEM was first compared with TPD method at 745 nm for diffuse reflectance measurements of optical phantoms, *ex vivo* tissues, as well as *in vivo* tissues. The DEM was then compared against SPD methods at 745 nm for the extraction of optical properties of the same samples, whereas the ground truth was obtained from the TPD method. In the rest of this work, TPD method with 5 spatial frequencies was also used as ground truth for optical property and chromophore concentration measurements [21], [22].

While the publicly available version produced notable artifacts when directly applied to the collected images in this study, the SPD models were re-trained following the network structure and procedures described in Aguénounon et al. with the corresponding published code [23]. Specifically, four CNNs for SPD were trained for the four *f*_*x*_ combinations as in Aguénounon et al., i.e., [0, 0.1], [0, 0.2], [0, 0.3], and [0, 0.4] mm^-1^. We note that the developed deep learning models were used for demodulation, while the diffuse reflectance and optical properties were extracted following standard procedures in the field. Additionally, the training of the DEM and SPD were conducted using the same dataset.

Moreover, to validate the generalization capability to out-of-sample wavelengths, the proposed DEM and the SPD methods were also used to extract optical properties for phantoms, *ex vivo* tissues, and *in vivo* tissues, at 655 nm and 850 nm wavelengths, and compared against ground truth values obtained from TPD method.

With optical properties of the three wavelengths, chromophore concentrations can be estimated for oxyhemoglobin and deoxyhemoglobin using Beer’s law. The proposed DEM was further compared with SPD methods for chromophore concentration extraction, whereas the ground truth was obtained from conventional TPD method.

### E. Cuff Occlusion Experiment

The cuff occlusion study was conducted in accordance with the institutionally approved protocol. To demonstrate longitudinal measurement of tissue hemodynamics, the hand of the subject was measured with the proposed method during a cuff occlusion study. The measurement was conducted for 8.5 minutes using [655, 745, 850] nm wavelengths, and repeated every 2.2 seconds. After approximately 3 minutes of baseline measurement, the pressure in the cuff (applied to the upper arm) was rapidly increased to ∼200 mm Hg and lasted for 2 minutes. Then, the cuff was released, and measurement continued for an additional 3.5 minutes. The hemoglobin concentrations were extracted with optical absorption of the three wavelengths using Beer’s law.

## IV. Result

### A. Diffuse Reflectance

The proposed method was first evaluated on the test set (n=75) for diffuse reflectance measurement at 745 nm wavelength. Fig. 3(a), (b), and (c) show the diffuse reflectance images of optical phantoms, *ex vivo* tissue (pork liver), and *in vivo* tissue, respectively. Visually, the diffuse reflectance maps obtained from the proposed DEM appear almost identical with the ground truth images from TPD method. Quantitatively, the average diffuse reflectance values for all test samples were calculated using a region-of-interest (ROI) of 22500 pixels (e.g., a square region of 150 × 150 pixels) at the center of the FOV, as shown in Fig. 4(a). The average values of different spatial frequencies obtained with SPD and DEM methods were respectively compared with those from the TPD (ground truth), as shown in Fig. 4(b) and (c), respectively. The choice of spatial frequencies for SPD were with reference to Aguénounon et al. [23]. The choice of spatial frequencies for DEM were with reference to the TPD [21], [22]. The overall errors of SPD and DEM methods were 0.65 ± 2.3% and 0. 85 ± 4.0%, respectively, indicating accurate measurement of diffuse reflectance values with both methods. The diffuse reflectance error from DEM is slightly larger than that from SPD, potentially due to the fact that the DEM simultaneously extracts much more spatial frequencies (2.5×) compared to SPD. Interestingly, as will be demonstrated below, the extracted multiple spatial frequencies by DEM, when combined, leads to significantly more accurate estimation of optical properties compared to the SPD.

**Fig. 3.**
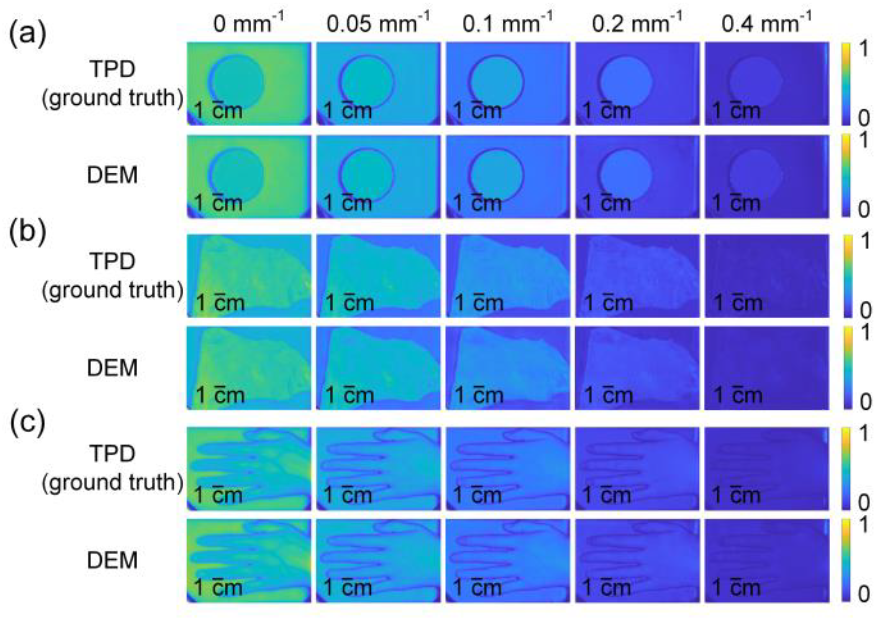
Diffuse reflectance maps of TPD (ground truth) and DEM for (a) optical phantom, (b) *ex vivo* tissue (pork liver), and (c) *in vivo* tissue (human hand). From left to right, the spatial frequencies are 0 mm^−1^, 0.05 mm^−1^, 0.1 mm ^−1^, 0.2 mm ^−1^, and 0.4 mm ^−1^, respectively.

**Fig. 4.**
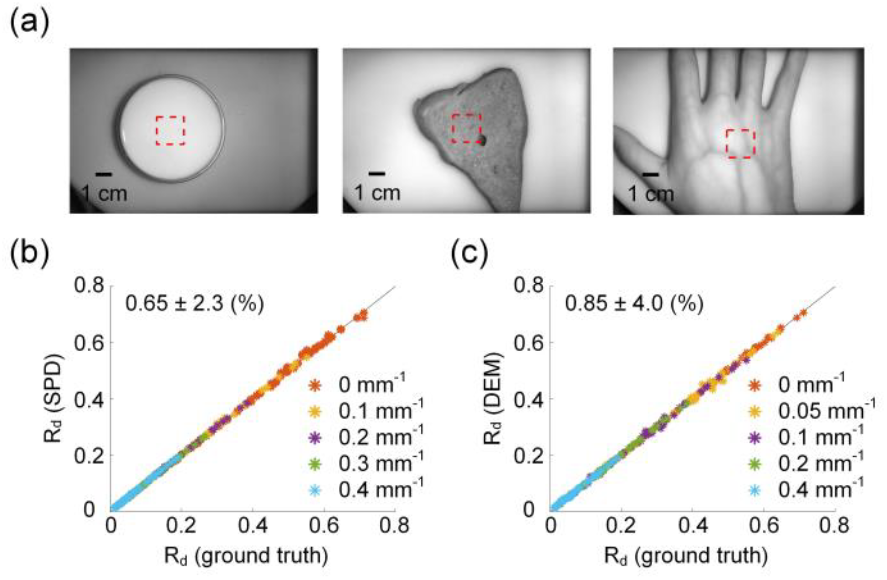
Comparison of diffuse reflectance from SPD and DEM against ground truth. (a) Representative ROIs. (b) Comparison of diffuse reflectance from SPD against TPD (ground truth) for all test samples at 745 nm wavelength. (c) Comparison of diffuse reflectance from DEM against TPD (ground truth) for all test samples at 745 nm wavelength.

### B. Optical Properties at Training Wavelength

Fig. 5 shows extracted OP maps with TPD (ground truth), SPD, and DEM, respectively, for optical phantom, *ex vivo* tissue (pork stomach), and *in vivo* tissue. Fig. 5(a) shows the OP maps of a homogeneous phantom, where imaging artifacts can be observed in the SPD maps. Moreover, Fig. 5(b) and Fig. 5(c) show OP maps of *ex vivo* pork tissue and *in vivo* human hand tissue, respectively. Compared to the ground truth, SPD methods give noticeable artifacts with different spatial frequency combinations. In contrast, the OP maps from DEM show better agreement with the ground truth images, suggesting improved measurement efficacy compared to the SPD.

**Fig. 5.**
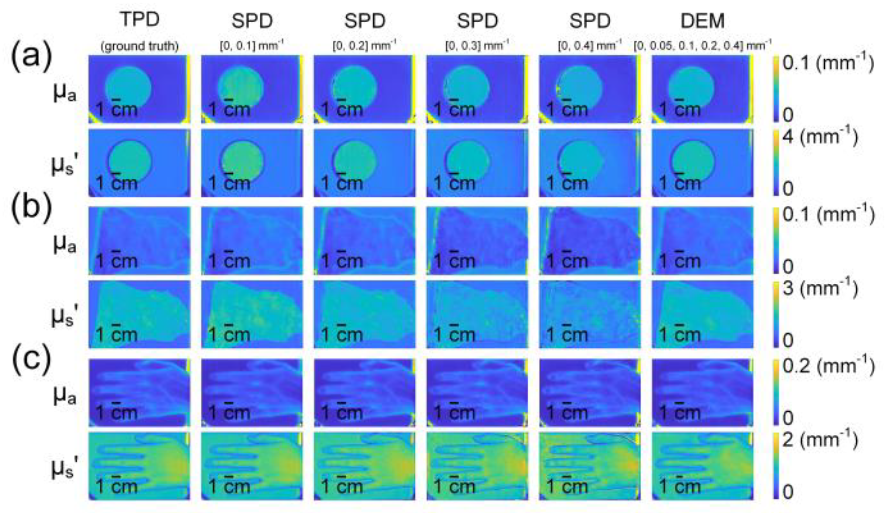
OP maps from TPD (ground truth), SPD (with different spatial frequency combinations), and DEM for (a) optical phantom, (b) *ex vivo* tissue (pork stomach), and (c) *in vivo* human hand tissue.

To quantify measurement discrepancies, we used the same ROIs as in Fig. 4 to compare OP measurements from different methods. Fig. 6(a) compares OP measurements of SPD against expected values obtained from TPD method, with different colors representing results from varied spatial frequency combinations. The overall measurement errors of SPD with different *f*_*x*_ combinations were 2.5 ± 15% and -1.2 ± 11% for optical absorption and reduced scattering, respectively. In contrast, Fig. 6(b) compares OP measurements of the proposed DEM method with expected values. The measurement errors were 0.83 ± 5.0% and 0.40 ± 1.9% for absorption and reduced scattering, respectively. The experimental results demonstrate that the proposed method has significantly improved accuracy for optical property measurements compared to SPD method.

**Fig. 6.**
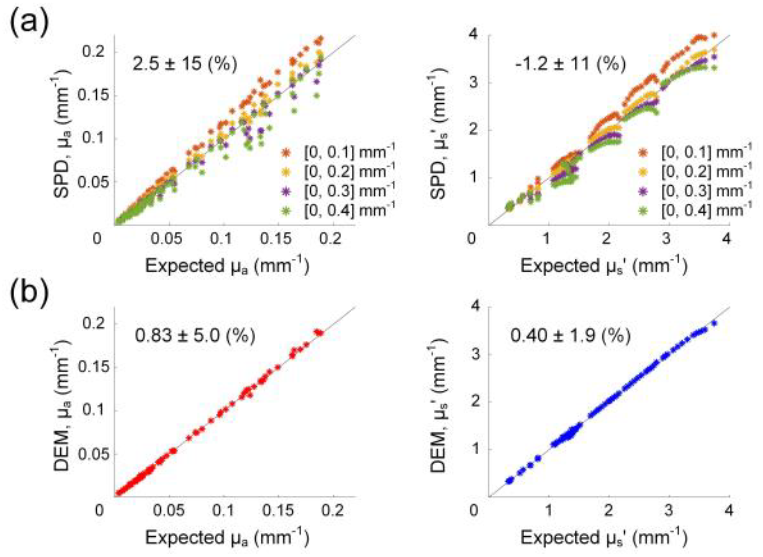
Comparison of optical property measurements. (a) SPD with different *f*_*x*_ combinations. The overall errors of OP extraction were 2.5 ± 15% and -1.2 ± 11% for absorption and reduce scattering, respectively. (b) Proposed DEM method. The errors of OP extraction were 0.83 ±5.0% and 0.40 ± 1.9% for absorption and reduce scattering, respectively, demonstrating significantly improved accuracy compared to SPD method.

### C. Optical Properties at Non-training Wavelengths

In practical scenarios, the developed model might need to be applied to wavelengths that were not included in training. To evaluate the generalization capability, the trained models from both SPD and the proposed method were directly applied to measurement data from the other two wavelengths, i.e., 655 nm and 850 nm, without any retraining of the deep learning models. Fig. 7(a) shows OP measurements of SPD models (with different spatial frequency combinations as in Fig. 6) against expected values obtained from the TPD at two non-training wavelengths. The measurement errors were -1.3 ± 19% and - 1.6 ± 13% for optical absorption and reduced scattering, respectively. In contrast, Fig. 7(b) compares OP measurements of the proposed DEM method with expected values. The measurement errors were -0.47 ± 19% and 1. 1 ± 5. 7% for optical absorption and reduced scattering, respectively. The experimental results demonstrate that the proposed method has improved accuracy for OP measurements compared to SPD methods at out-of-sample wavelengths not included in training.

**Fig. 7.**
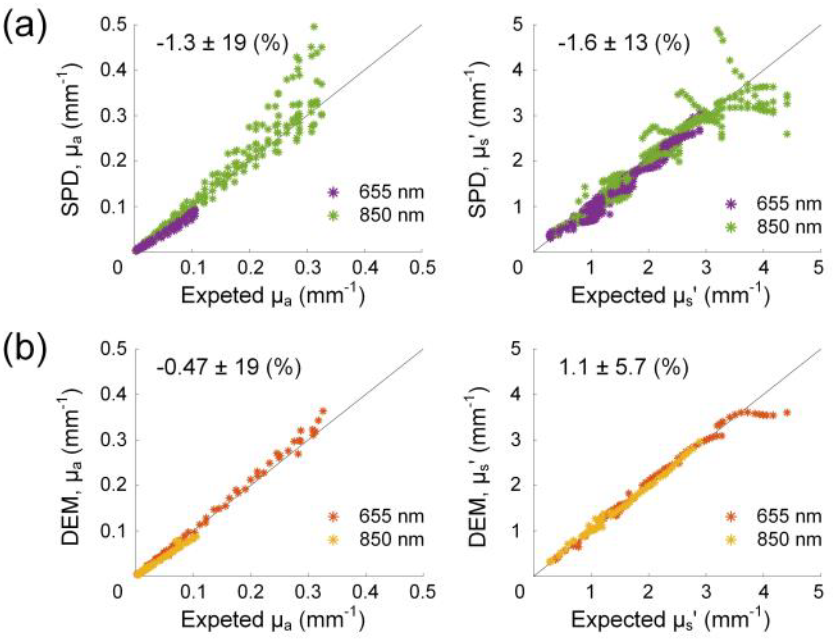
Comparison of OP measurements at non-training wavelengths. (a) SPD method. (b) The proposed DEM method.

### D. Chromophore Concentrations and Tissue Oxygenation

Fig. 8 shows hemoglobin concentration and tissue oxygenation maps obtained from the same subject using TPD, SPD of different *f*_*x*_ combinations, and DEM, respectively. Fig. 8(a)-(d) compare oxyhemoglobin concentration (HbO2), deoxyhemoglobin concentration (Hb), total hemoglobin concentration (THb), and tissue oxygenation (StO2), respectively. While TPD and DEM both present wide-field measurements of *in vivo* tissue with little artifact, the images from SPD have severe artifacts, especially for oxyhemoglobin maps and tissue oxygenation maps.

**Fig. 8.**
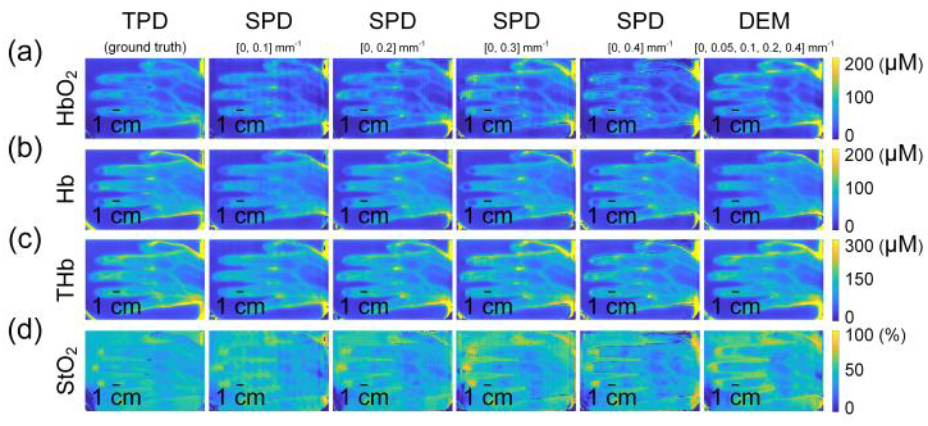
Chromophore concentration and tissue oxygenation maps obtained with different methods. (a) Oxyhemoglobin concentration. (b) Deoxyhemoglobin concentration. (c) Total hemoglobin concentration. (d) Tissue oxygenation. From left to right: TPD, SPD with different *f*_*x*_ combinations, and DEM.

To quantitatively compare the discrepancies of chromophore concentrations and tissue oxygenation, the values obtained with TPD were used as ground truth, and the same ROIs as in Fig. 4 were used for calculation (n=8 from the test set). As shown in Table 1, the errors with DEM were generally lower compared to those of SPD with different *f*_*x*_ combinations.

**TABLE I.** Quantitative Comparison for Percent Errors (%) of Chromophore Concentrations and Tissue Oxygenation With Different Methods

|  | SPD<br>[0, 0.1]<br>mm<sup>-1</sup> | SPD<br>[0, 0.2]<br>mm<sup>-1</sup> | SPD<br>[0, 0.3]<br>mm<sup>-1</sup> | SPD<br>[0, 0.4]<br>mm<sup>-1</sup> | DEM |
| --- | --- | --- | --- | --- | --- |
| HbO<sub>2</sub> | 16 ± 6.6 | 20 ± 7.4 | 22 ± 11 | 21 ± 6.7 | 13 ± 7.2 |
| Hb | 3.4 ± 6.9 | 0.48 ± 2.8 | 2.5 ± 3.0 | 8.3 ± 3.8 | 3.0 ± 2.6 |
| THb | 9.4 ± 4.9 | 9.4 ± 2.8 | 12 ± 4.4 | 14 ± 3.4 | 7.6 ± 2.8 |
| StO<sub>2</sub> | 7.6 ± 6.3 | 12 ± 6.7 | 12 ± 10 | 8.9 ± 6.4 | 6.1 ± 5.8 |

### E. Cuff Occlusion Experiment

Fig. 9 shows the chromophore concentration maps from the final frame of the recorded video for the cuff occlusion including the time series of hemoglobin changes calculated from the ROI (red-dashed boxes). The video was sped up 45× for the ease of viewing (Visualization 1). The experimental results demonstrate that the hemodynamics of *in vivo* human tissue can be longitudinally monitored with the proposed method.

**Fig. 9.**
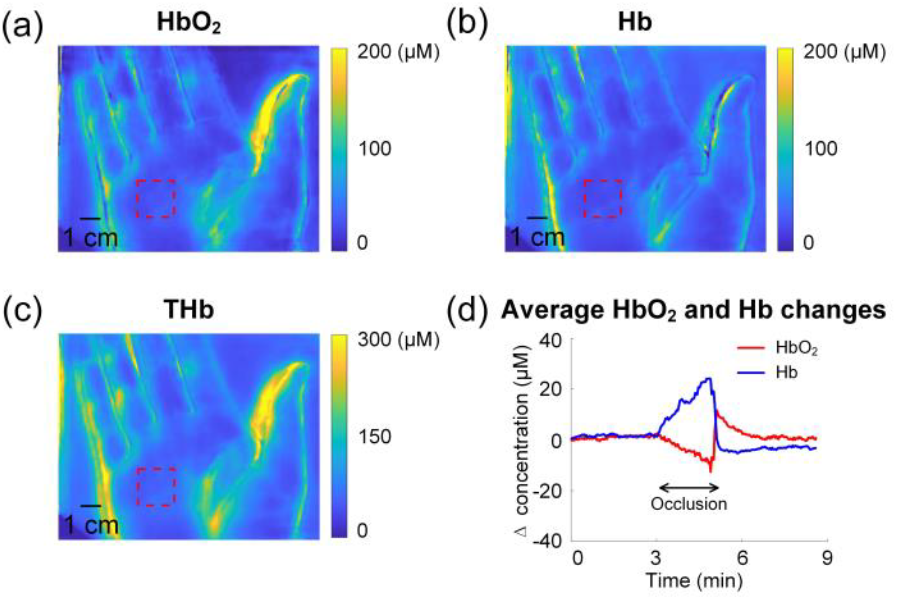
Cuff occlusion measurements. (a) Oxyhemoglobin concentration. (b) Deoxyhemoglobin concentration. (c) Total hemoglobin concentration. (d) Changes in hemoglobin concentrations during the cuff occlusion experiment.

## V. Discussion

We have developed and validated DEM as a new model for high-speed and accurate mapping of tissue optical properties in a single snapshot. This is also enabled through frequency multiplexing, i.e., the use of specifically designed illumination pattern that contains five angularly rotated *f*_*x*_ patterns, which allows access to multiple spatial frequencies in a single reflectance image. Intuitively, one would attempt the extraction of *f*_*x*_ information (i.e., demodulation) through filtering and inverse transformation in the Fourier domain. However, that only worked on limited phantom measurements, and would produce severe artifacts on tissue measurements. Therefore, we proposed the deep ensemble model which can extract demodulated images of each spatial frequency without producing artifacts. The demodulated images of different spatial frequencies were then used to extract optical properties. Through validation on an array of optical phantoms, *ex vivo* tissues, and *in vivo* tissues, we demonstrated that compared to the TPD method (ground truth), the proposed method gave accurate measurement values on diffuse reflectance as well as optical absorption and reduced scattering. We further validated that while trained with measurements of a single wavelength, the proposed DEM can be directly applied to wavelengths not included in the model training and can also produce accurate optical property values. We then demonstrated the DEM for quantitative extraction of chromophore concentrations and tissue oxygenation using three measurement wavelengths. Finally, to highlight the single snapshot imaging capability for tissue optical properties enabled by the proposed method, we demonstrated longitudinal monitoring of *in vivo* human tissue hemodynamics with quantitative mapping of chromophore concentrations through a cuff occlusion experiment.

DEM enjoys a number of advantages over methods currently used for mapping tissue optical properties. Conventional TPD method can also quantify optical properties with high accuracy, but it requires a total number of 14 measurement images. In contrast, the proposed DEM only requires a single measurement image, and is 14× faster compared to the TPD. Additionally, Nadeau et al. developed a multifrequency synthesis and extraction method as an alternative of the TPD. However, it required 7 uniformly phase-offset square wave patterns for the extraction of DC and three harmonic components for *in vivo* measurements [33]. Moreover, SPD methods can also quantify optical properties with a single measurement image, but are limited to 2 spatial frequencies and larger measurement uncertainties. In contrast, the proposed DEM method can extract OPs using 5 spatial frequencies with a single snapshot image, and has considerably lower measurement uncertainties. Additionally, compared to the SPD methods, the proposed method has lower quantitative errors as well as significantly reduced image artifacts for the extraction of chromophore concentrations and tissue oxygenation, which can be particularly helpful in surgical monitoring applications [34]. It is also important to note that in addition to biological tissues, the proposed method can be applied to quantitative mapping of optical properties for other strongly scattering media such as turbid flow, which may have substantial impact for fluid dynamics [35].

Going forward, there are several ways to further improve and expand the capabilities of the proposed DEM. For example, the accuracy of the deep ensemble model can potentially be improved by using training data from more samples and wavelengths [36], [37]. The ensemble framework can also be adopted for other *f*_*x*_ combinations. For instance, more *f*_*x*_s may be used to further reduce measurement uncertainties [21]. Additionally, the proposed pattern can potentially be printed on a transparency film for projection without the use of DMD, which may significantly reduce the cost in system hardware [35], [38]. Furthermore, the proposed method could be particularly useful for visible wavelengths, where OP values are typically high, and measurement uncertainties could be relatively large. Also, with instant measurement of multiple spatial frequencies, one can potentially achieve “snapshot tomography”, given that different *f*_*x*_s have different depth sensitivity [39]. Finally, the deep learning model could be trained to process the measurement image directly to tissue optical properties, further enabling real-time feedback during clinical applications [40].

## VI. Conclusion

In summary, this work introduced a deep ensemble model that can extract optical properties with high accuracy from a single snapshot, increasing the measurement speed by 14× compared to conventional TPD. With validation on optical phantoms, *ex vivo* tissues, and *in vivo* tissues, the proposed DEM method has higher measurement accuracy and reduced image artifacts compared to the state-of-the-art SPD methods. The deep ensemble model for demodulation, although trained on a single wavelength, can be directly applied to other wavelengths, and then used for the estimation of chromophore concentrations as well as tissue oxygenation. Taken together, the proposed method paves the way for high-speed, accurate, and wide-field monitoring of tissue optical properties, which can be helpful for a variety of biomedical applications including small animal imaging, burn wound assessment and surgical monitoring.

## Supporting information

hemodynamics monitoring

## Acknowledgment

The authors gratefully thank Dr. Darren Roblyer at Boston University and Dr. Yan Xu at Beihang University for helpful discussions.

